# Transitions in brain-network level information processing dynamics are driven by alterations in neural gain

**DOI:** 10.1101/581538

**Authors:** Mike Li, Yinuo Han, Matthew J. Aburn, Michael Breakspear, Russell A. Poldrack, James M. Shine, Joseph T. Lizier

**Affiliations:** Centre for Complex Systems, The University of Sydney, Sydney, Australia; Brain and Mind Centre, The University of Sydney, Sydney. Australia; Complex Systems Research Group, Faculty of Engineering and IT, The University of Sydney, Sydney, Australia; QIMR Berghofer Medical Research Institute, Queensland, Australia; Department of Psychology, Stanford University, USA

## Abstract

A key component of the flexibility and complexity of the brain is its ability to dynamically adapt its functional network structure between integrated and segregated brain states depending on the demands of different cognitive tasks. Integrated states are prevalent when performing tasks of high complexity, such as maintaining items in working memory, consistent with models of a global workspace architecture. Recent work has suggested that the balance between integration and segregation is under the control of ascending neuromodulatory systems, such as the noradrenergic system. In a previous large-scale nonlinear oscillator model of neuronal network dynamics, we showed that manipulating neural gain led to a ‘critical’ transition in phase synchrony that was associated with a shift from segregated to integrated topology, thus confirming our original prediction. In this study, we advance these results by demonstrating that the gain-mediated phase transition is characterized by a shift in the underlying dynamics of neural information processing. Specifically, the dynamics of the subcritical (segregated) regime are dominated by information storage, whereas the supercritical (integrated) regime is associated with increased information transfer (measured via transfer entropy). Operating near to the critical regime with respect to modulating neural gain would thus appear to provide computational advantages, offering flexibility in the information processing that can be performed with only subtle changes in gain control. Our results thus link studies of whole-brain network topology and the ascending arousal system with information processing dynamics, and suggest that the constraints imposed by the ascending arousal system constrain low-dimensional modes of information processing within the brain.

**Author summary:** Higher brain function relies on a dynamic balance between functional integration and segregation. Previous work has shown that this balance is mediated in part by alterations in neural gain, which are thought to relate to projections from ascending neuromodulatory nuclei, such as the locus coeruleus. Here, we extend this work by demonstrating that the modulation of neural gain alters the information processing dynamics of the neural components of a biophysical neural model. Specifically, we find that low levels of neural gain are characterized by high Active Information Storage, whereas higher levels of neural gain are associated with an increase in inter-regional Transfer Entropy. Our results suggest that the modulation of neural gain via the ascending arousal system may fundamentally alter the information processing mode of the brain, which in turn has important implications for understanding the biophysical basis of cognition.

## Introduction

Although there is a long history relating individual brain regions to specific and specialized functions, regions in isolation cannot perform meaningful physiological or cognitive processes [1]. Instead, starting at a lower scale, a few prominent features of the brain’s basic mechanisms stand out. Firstly, neurons exist in vast numbers, each acting as an individual element with a similar set of rules. Secondly, the response of individual neurons to stimuli are far from linear – small changes in the surrounding milieu can lead to abrupt changes in neural dynamics [2]. Thirdly, all neurons interact with other neurons through synapses, and hence form a network that spans the central nervous system [3]. Furthermore, this structural backbone supports coherence of physiological activity at larger scales, giving rise to distributed functional networks [4]. Therefore, in every regard, the brain is a complex system whose computational power stems from the emergent properties of coordinated interactions between its components [5, 6]. Understanding how the topology *and* dynamics of these networks give rise to its function is one of the most central questions that computational neuroscience aims to address.

From comparing a range of physical and mathematical systems, it is known that complex systems can exist in multiple distinct phases. For instance, groups of water molecules can exist as a solid, liquid or gas, depending on the surrounding temperature and pressure. By altering one or more tuning parameters (e.g. temperature in the water example), the system can cross clearly defined critical boundaries in the parameter space. These critical transitions are typically abrupt and often associated with qualitative shifts in the function of a system (e.g. consider the stark differences between ice and liquid water). They are often of great interest due to their ubiquity and the implications for systemic flexibility [8].

Empirical observations in neural cultures, EEG and fMRI recordings provide evidence that the brain operates near criticality [8–16] – one form of which is a transition between two distinct states in the functional network topology [17] (see Fig. 1c). At one extreme, different regions of the brain are highly segregated, and each region prioritizes communication within its local topological neighbourhood. At the other pole, the whole brain becomes highly integrated, and cross-regional communication becomes far more prominent. Experimentally, the resting brain is found to trace a trajectory between the two states, and can transition abruptly into the highly integrated state when the subject is presented with a cognitively challenging task [18].

As described in Methods regarding [7], this transition across a critical boundary can be achieved in a neural mass model by tuning two parameters: the neural gain (*σ*) and excitability (γ). Fig. 1a shows how the gain parameter increases each region’s signal-to-noise ratio by altering the shape of the input-output curve, while the excitability scales the magnitude of the response. Biologically, the modulation in precision and responsivity of targeted neurons is mediated by ascending noradrenergic projections from regions such as the locus coeruleus [7, 19]. This tuning of neural gain mediated by noradrenaline can significantly alter the functional network topology of the brain, as characterized by graph theoretical parameters, such as the mean participation coefficient, and temporal measures, such as the phase synchrony order parameter. In Fig. 1b, the sharp transition in mean synchrony requires only a small change in neural gain, and is an illustration of the critical behaviour of the network in that region of parameter space.

**Fig 1.**
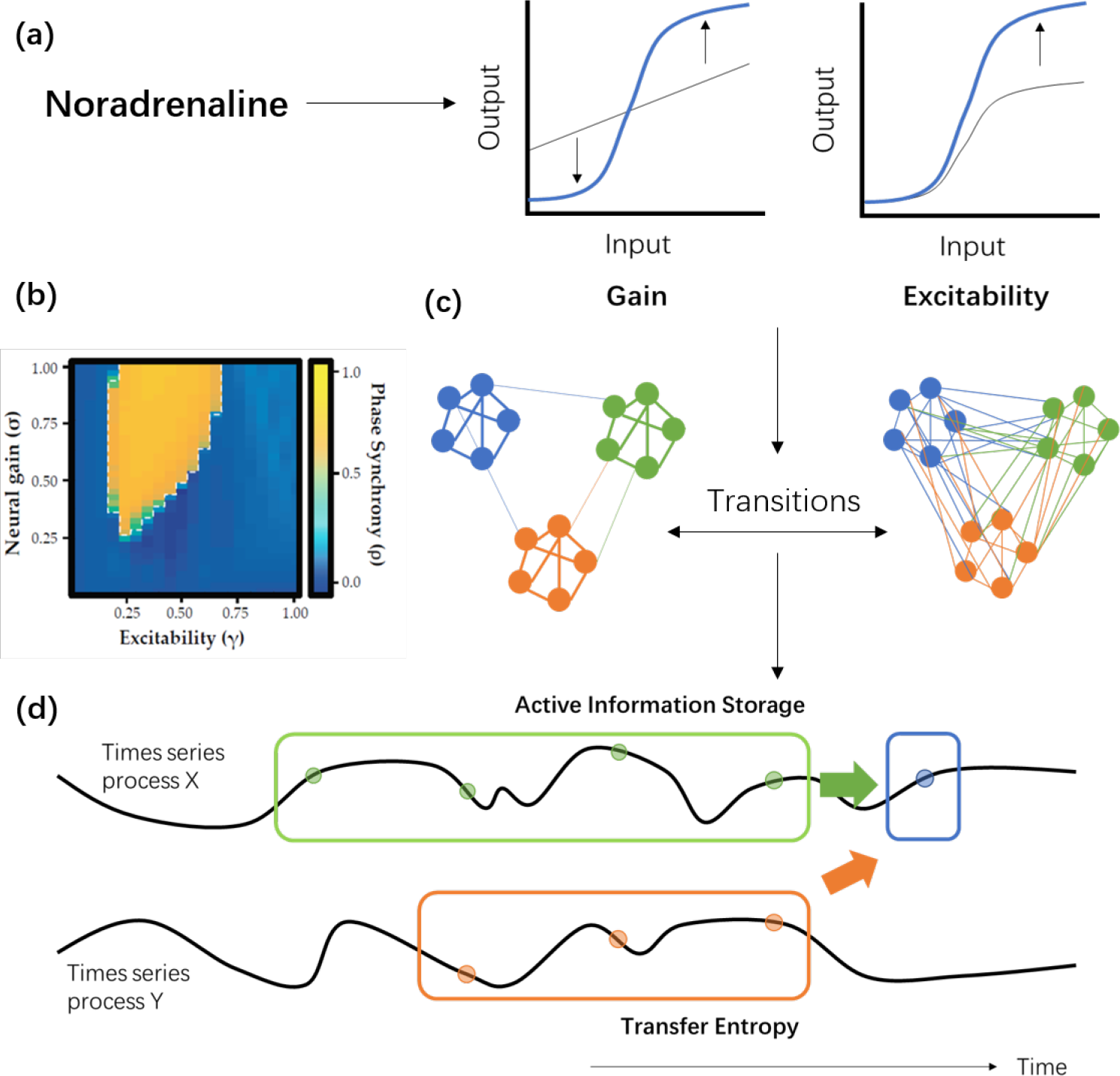
Schematic diagram showing how noradrenaline may potentially affect the information processing structure of the brain. (a) Noradrenaline plays an important role in tuning the neural gain of targeted neurons. The effect of neural gain and excitability, the two tuning parameters being varied in our neural mass model, on the response of individual neurons to stimuli are shown schematically. (b) Previous results from [7] (reproduced under Creative Commons Attribution License CC BY 4.0) showing that varying neural gain and excitability may cause abrupt changes in the mean phase synchrony of the brain from modelled fMRI BOLD recordings, implying the existence of a critical boundary between a segregated phase (low phase synchrony) and an integrated phase (high phase synchrony) in the brain. (c) Schematic diagram of brain regions in the segregated and integrated phases, and how changing neural gain and excitability may lead to transitions between the two. (d) Schematic diagram of the concept of active information storage and transfer entropy, and how they may be affected by phase transitions. Qualitatively, active information storage (green arrow) describes information on the next instance *X*_*n*+1_ (blue sample) of a time series *X* provided by its own history (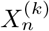, green samples), whereas transfer entropy (orange arrow) describes that provided by the past (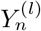, orange samples) of another time series *Y* in the context of the target’s history. See further details on these measures in Methods.

A question then arises: why might it be favourable for the brain to be in a near or quasi-critical state in the first place? Specifically, are there computational advantages accompanying this structure? Many have proposed that this signature may reflect an evolutionary optimisation, allowing for both an effective balance between long- and short-range interactions between neural regions, as well as rapid transitions between segregated and integrated states [20, 21]. For instance, a highly segregated brain cannot communicate effectively to share information across different sub-networks, while a highly integrated state results in homogeneity in the flow of signals and a reduction in meaningful interactions, as in the case of epilepsy [18]. Hence, an optimal state for the brain is likely a flexible balance between the two extremes. Indeed, systems poised near criticality are well-known to exhibit other distinct characteristics which could be usefully exploited in the brain, such as increased autocorrelation times and variance [22–24], increased coupling across the system [25, 26] and maximal sensitivity to tuning parameters [27].

Prompted by early conjecture [28], there is evidence from neural recordings and studies of other complex systems that phase transitions are often related to changes in the *information processing* structure of a system. Shew et al. [10] demonstrated maximal information capacity (via entropy) and sharing of information (via mutual information) near critical transitions in dynamics of neural cultures, with the transitions investigated by manipulating excitation-inhibition ratios. Although the study referred to the latter measure as “information transmission”, the mutual information remains a measure of statically shared information, and information transmission and processing in general are more appropriately modelled with measures of dynamic state updates [29]. Such measures have provided more direct evidence of changes in information processing structure associated with phase transitions (in the brain, in preliminary results of Priesemann et al. [30], and) in other complex systems. In artificial recurrent neural networks for example, both information transfer and storage were observed to be maximized close to a critical phase transition (with respect to perturbation propagation in reservoir dynamics) [31], suggesting that these intrinsic information processing advantages underpinned the known [32] higher performance of similar networks near the critical point on various computational tasks. These changes in information processing structure can also explain some of the aforementioned characteristics near the critical point, such as increased autocorrelation times (as a result of elevated information storage) and coupling (as a result of increased information transfer). Similar results are seen in the well-known phase transition with respect to temperature in the Ising model, with information transfer maximized near the critical regime [33]. Furthermore, the dynamics of Boolean networks (models for gene regulatory networks [34]) exhibiting order-chaos phase transitions are dominated by information storage in the ordered low-activity phase, and information transfer in the high-activity chaotic phase [35, 36]. At the critical regime, networks exhibit a balance by combining relatively strong capabilities of both information storage and transfer. This transition in Boolean networks can be triggered either by directly altering the level of activity in the dynamical rules of the nodes, or by sweeping the randomness in network structure starting with a regular lattice network (low-activity) through small-world [37] and onto random structure (high-activity). The dynamics of the brain, being a highly analogous system to these exhibiting phase transitions between functional segregation and integration, may exhibit similar patterns in information storage and transfer capabilities near the critical regime, and on both sides of the critical boundary. Hence, we aim to examine the quantitative changes in the information processing properties of the brain under the framework of information theory.

From a Shannon information-theoretic perspective, we measure information as the reduction in uncertainty about an event with an unknown outcome [38]. For a given time series process within a larger system, such as the blood oxygen level dependent (BOLD) data for a single voxel in the brain (i.e. the smallest identifiable region in an fMRI scan), the sources of information regarding the next event in the process include the history of the time series of the process itself, and the history of other processes in the system as inputs, such as the time series of other voxels. Here, within the framework of information dynamics [29] we model the amount of information storage as that provided from within a time series process using the active information storage (AIS) [39]. We model information transfer as that provided by another source to a target process, in the context of the target past, using transfer entropy (TE) [40]. Fig. 1d provides a simple illustration of this concept.

By using computational modelling to examine the behaviour of these two information theoretic measures across a parameter space of varying values of neural gain and excitability, we aim to address two main questions: firstly, are there differences in the information processing structure as a function of neural gain alteration? And secondly, does a quasi-critical state provide computational information processing benefits? Given the properties of the previously determined topological measures (Fig. 1b) and how they relate to previous results on information processing around critical regimes, we predicted differential information processing structures across the parameter space of gain and excitability, and hypothesized that: i. the active information storage across the system should be maximized in the subcritical region before the critical boundary, whereas ii. the transfer entropy would be maximized after the boundary in the supercritical region, and iii. that storage and transfer should be relatively balanced at the critical transition. A change in information processing near criticality may allow for rapid alterations in the balance between states dominated by information storage in the subcritical phase and information transfer in the supercritical phase, hence providing flexibility for the dynamical structure of the brain to quickly adapt to and complete a wide range of tasks.

## Results

Regional time series of neuronal dynamics were generated by a 2-dimensional neural oscillator model with stochastic noise [41] built on top of a weighted, directed white matter connectome [42], simulated with the Virtual Brain toolbox [43]. The properties of inter-regional coupling were systematically adjusted using the parameters for gain (*σ*) and excitability (γ) (see Methods for more details).

In contrast to the previous study which used the same underlying generative model [7], we did not transform the raw data into a simulated BOLD signal. Instead, information theoretic measures were calculated directly on each region’s average membrane voltage (which is monotonically related to the neural firing rate) – sampled at a rate of 2 kHz. This allowed us to construct a model of information processing that was more closely linked with the underlying dynamics of the neural system. For each point in the two dimensional *σ*-γ parameter space, information storage was calculated for each region from its own generated time series process. Similarly, information transfer was calculated for each directed pair of regions from their own generated time series processes at each point in the *σ*-γ space. The fast sampling rate obliged us to apply the information theoretic measures treating the data as arising from a continuous-time process (see Methods).

### Information storage peaks in the subcritical region at intermediate γ

The active information storage of a process measures the extent to which one can model the next sample of a time series as being computed from (a time-delay phase space embedding of) its past history [39] (see Methods). High active information storage implies that the past states of a process are strongly predictive of the next observation. For this experiment, we measured the active memory utilization rate (AM rate), which is a formulation of active information storage suitable for continuous time processes [44] (see Methods).

Fig. 2a plots the active memory rate (averaged over all regions) with respect to the *σ*-γ space. Active memory rate peaks at what was previously identified as the subcritical or segregated regime [7] (compare to regime identification in Fig. 1b). We also see two types of transitions that divide the space up into four qualitative regions. One transition occurs over variations in γ: the highest information storage occurs at intermediate values of γ, with a sharp dropoff on either side. Within this band of intermediate γ there is also a transition in *σ*, with the highest information storage occurring at small values of *σ*, again with a sharp dropoff across the critical transition.

This correspondence with the previously identified regions can be observed more clearly from Fig. 2b and Fig. 2c, which plots the average values across the *σ* and γ boundaries, respectively, based on the synchronization order parameter of the model from [7]. A qualitative change is observed at these boundaries, where the active memory rate is highest (and peaking) in the subcritical regime with respect to *σ*, whilst still exhibiting sharp transitions to higher values in the supercritical regime with respect to γ.

### Correlation of information storage to motif counts suggests distributed memory in supercritical regime

Information storage in time-series dynamics of network-embedded processes is known to be supported by certain loop motif structures within directed structural networks, such as low order feed-forward and feedback loops, and under certain types of simple dynamics an exact relationship can be derived [45]. For the dynamics used with this model, we do not know the exact relationship, but we can still approximate the relative local network support provided for information to be stored in the time-series process at a particular region. This allows us to check how the influence on storage of these relevant structural motifs changes with the parameter space. The local network support is computed as a linear weighted path sum of different types of motifs (of length larger than 1) originating in each region (see some examples in Fig. 2d, and Methods for details). We correlate this local network support for each region with its active memory rate, to examine whether and where in the phase space of dynamics these motifs are indeed strongly supporting information storage.

Fig. 2e shows a high correlation between the active memory rate and local network support in the supercritical phase. This fits with established results of [45] that assume a noise-driven system. This can be contrasted with the correlation of information storage to normalized within-region synaptic connection weight (self loop weights *A*_*ii*_ in (3) in Methods) in Fig. 2f, which peaks in the high γ subcritical regime. This suggests that primarily synaptic connections internal to a region support information storage in the segregated dynamics of the sub-critical regime, whilst during the supercritical regime we observe the engagement of network effects via longer motifs to support more distributed memory in the integrated dynamics. Interestingly, we note that the subcritical regime with strongest information storage (intermediate γ, low *σ*) appears to have support from both within-region synaptic connections and longer motifs.

**Fig 2.**
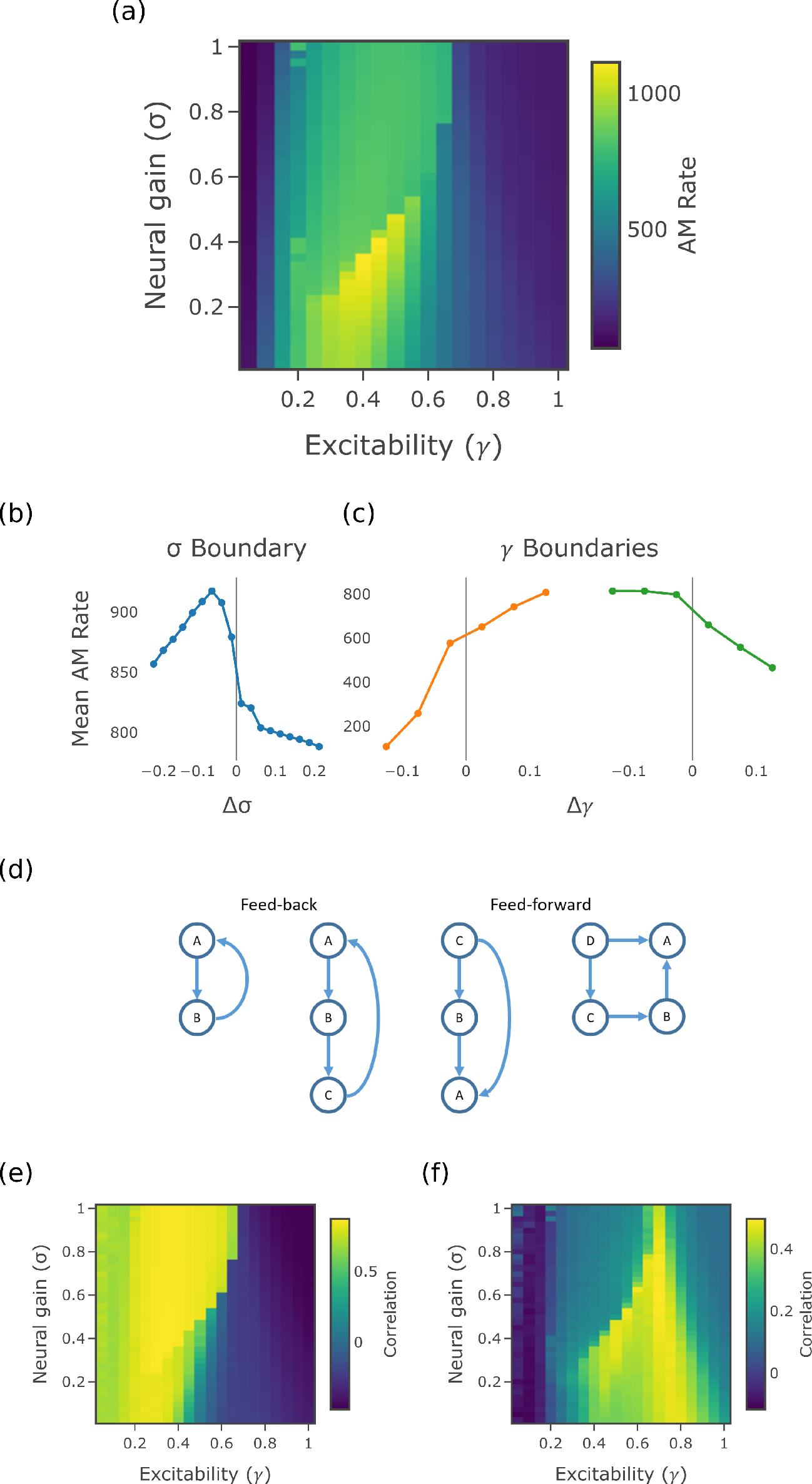
Measures of information storage. (a) Active memory utilization rate. (b) and (c) Mean active memory rate across *σ* and γ phase boundaries. (d) Network motifs supporting information storage in the dynamics of node A. (e) Correlation of AM rate to local network support (weighted motif counts). (f) Correlation of AM rate to normalized within-region synaptic connection weight.

### Information transfer is maximized in the supercritical region

Information transfer from one process to another is modelled by the transfer entropy [40] as the amount of information which a source provides about a target’s next state in the context of the target’s past (see Methods).

For this experiment on continuous-time processes, we measure the transfer entropy rate [46]. Unless otherwise stated, we constrain the information sources considering only those which are causal information contributors to the target (following [35, 47]); these are known from the directed structural connectome used in the simulation.

For each point in the parameter space of *σ* and γ, we calculate all the pairwise transfer entropy rates to targets from each of their causal parents. These values are averaged across all such directed pairs to give the mean (pairwise) transfer entropy rate across the network at each *σ* and γ pair. Fig. 3a shows a clear separation of the parameter space into the subcritical and supercritical regions (as defined in the previous work [7], see Fig. 3d and Fig. 3e), with information transfer occurring almost exclusively in the supercritical or integrated regime. An additional trend within the supercritical region can be seen, with TE rate rising as *σ* and γ increases.

Higher order terms for information transfer can also be calculated in addition to the pairwise components. Conditional transfer entropy [48, 49] (see Methods) adds the history of a third process or a collection of processes to be conditioned on in addition to the history of the target itself. For this experiment, we calculate the conditional transfer entropy of the causal source-target relationships, conditioned on all the other causal parents of the target (from the directed structural connectome), and average this across all directed causal pairs. In comparison to the pairwise TE, by conditioning on all the other causal parents the conditional TE captures only the unique information component which this source is able to provide that the others do not, and adds in synergistic or multivariate information about the target that it provides only in conjunction with the other sources. Furthermore, any information it holds about the target which is redundant with the other sources is removed.

The collective transfer entropy [49] (see Methods) captures the total transfer of information from a group of sources to a target. We examine the total information from the full set of causal parents to a particular target, and average this across all target processes. The collective TE captures all of the information provided by the sources about the target, whether that information is provided uniquely by any single source, or redundantly or synergistically by some set of them. Importantly, it does not “double count” information held redundantly across multiple sources.

Overall, the mean conditional TE rate in Fig. 3b and the mean collective TE rate in Fig. 3c show qualitatively similar trends to the pairwise TE rate in Fig. 3a, with the same strong mean transfer in the supercritical region and trend towards high γ and *σ* within that region, although this trend is even stronger with more of a skew towards high γ. From comparing the peak values of the different figures, it can be seen that substantial redundancies exist in the information held about the target between the different sources. That is, the conditional transfer entropy rate is an order of magnitude lower than the simple pairwise measure, suggesting that each causal parent is not able to provide much additional information beyond that already apparent from the other parents. The collective transfer entropy rate, on the other hand is an order of magnitude higher than the simple pairwise measure. This may, however, simply be due to the effect of adding up the transfer entropy rate from the different sources, and the collective transfer entropy divided by the number of sources is not actually higher than the pairwise measure. It is difficult to conclude from this whether there are network level effects that give rise to synergies beyond looking at each region in pairwise fashion (see Methods).

The behaviour of information transfer (Fig. 3) was observed to be complementary to the patterns of information storage (Fig. 2). This paints a picture of a distinct mode in the dynamics of information processing that switches abruptly as the system moves between the supercritical and subcritical phases. However, it should be noted that the values for transfer entropy rate are still smaller by one or two orders of magnitude, even in the collective case. This is partially due to the relative simplicity of our neural mass model, and also to the regularity of the oscillations which they produce.

**Fig 3.**
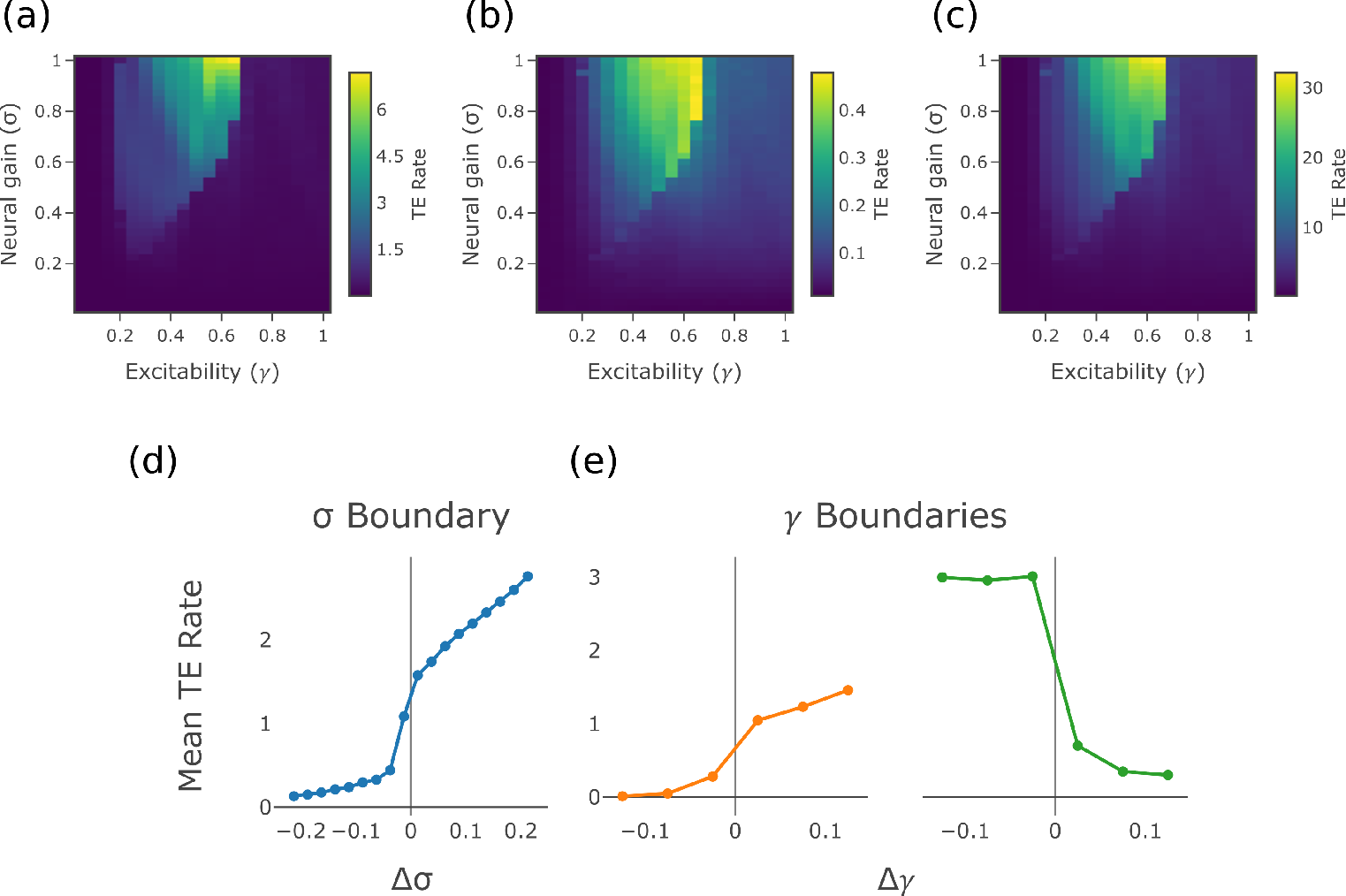
Measures of information transfer. (a) Average transfer entropy rate over causal edges (those connected source → target by the directed connectome). (b) Conditional transfer entropy rate over causal edges. (c) Collective transfer entropy rate of causal edges. (d) and (e) Mean TE rate across *σ* boundary and γ boundaries

### Correlation of information transfer with in-degree shows change in behaviour at the phase boundaries

Fig. 4a examines the correlation of the pairwise transfer entropy rate, averaged across all outgoing directed connections in the network for each source, with the in-degree of the source. This correlation is expected in general (having been observed in [50, 51]) because sources with higher in-degree have greater diversity of inputs, and so potentially more available information to transfer. The expected effect is observed in the supercritical phase, suggesting integration of the information from the different source inputs. Fig. 4b shows the correlation of conditional transfer entropy rate to source in-degree (again with the conditional TE rate measure averaged for all outgoing connections for each source), showing a similar pattern to Fig. 4a.

Fig. 4c shows the correlation of the collective transfer entropy rate for each target to target in-degree. The target in-degree is used instead of the source because one value of collective transfer entropy is produced for each target, while the contribution over all sources is combined. Because the collective TE rate captures the combined effect from all sources, it can be expected that this will increase with the in-degree of the target, and so the correlation should be quite strong. This is observed across the phase space, but is slightly weaker at the critical boundary (which on inspection appears to be due to larger non-linearities in the TE-degree relationship there).

**Fig 4.**
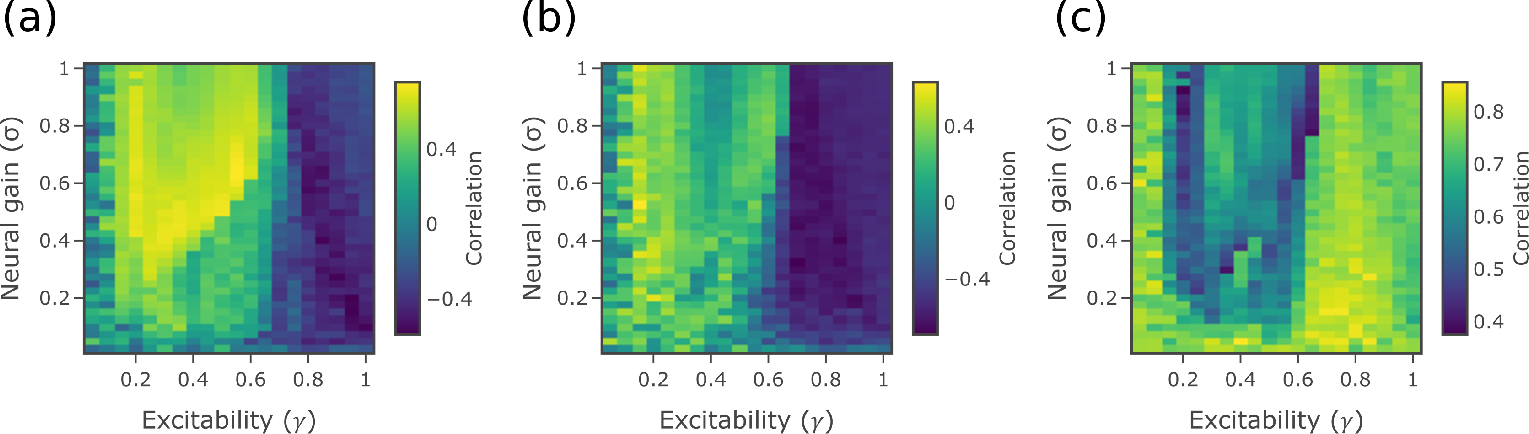
Correlations between information transfer and node degrees. (a) Correlation between TE rate and source in-degree. (b) Correlation between conditional TE rate and source in-degree. (c) Correlation between collective TE rate and target in-degree

### Inter-hemisphere information transfer is high in the supercritical region

The transfer entropy can be calculated solely for the causal edges which link regions between the two hemispheres. Only 38 links are inter-hemispheric (out of the total of 1494, not counting self loops). The weights of these connections are also relatively low, with an average weight of 1.07 (standard deviation of 0.86) compared to an average weight of 1.91 (standard deviation of 0.63) over all links, not counting self loops. Despite this, however, the average transfer entropy rate of these inter-hemisphere links in Fig. 5a follows the same pattern as the standard pairwise transfer entropy rate seen in Fig. 3a and the peak values are only 30% lower. This suggests that information transfer across hemispheres is significant, especially since Fig. 5a favours the high γ, high *σ* part of the supercritical region, which may help explain why this trend is seen in Fig. 3a.

Fig. 5b shows the outcome of a second test which again highlights the importance of inter-hemisphere information transfer in the supercritical phase. This figure looks beyond causal links to compare the proportion of total statistically significant pairwise transfer entropy which occurs between hemispheres. The transfer entropy rate is first calculated for all pairwise combinations of source and target (whether they are linked in the directed connectome or not), at each point in the parameter space. However, pairs which do not give a level of transfer entropy statistically different to zero are ignored (see Methods). The remaining pairs are used to calculate the proportion of total pairwise transfer entropies that are accounted for by inter-hemisphere transfer, for each point in the parameter space. This proportion is close to half in the supercritical phase, showing that there is a large indirect effect of the information transferred between hemispheres. Even though there are only a few causal links between hemispheres, the information transferred by these “long” links is novel and becomes redistributed within the hemisphere, underpinning the higher levels of integration observed in the supercritical phase.

## Discussion

Using an information theoretic decomposition, we extend previous work [7] by demonstrating that a gain-mediated phase transition in functional network topology is associated with a fundamental alteration in the information processing capacity of the whole brain network. Importantly, during this transition the underlying coupling strength and connectivity matrix are kept constant: the local dynamics are altered due to changes in the neural gain and excitability parameters, which then leads to changes in the effective connectivity (being a result of both local dynamics and large-scale structural connectivity). By modelling the distributed computation of the neural system in terms of information storage and information transfer, our results suggest that the shift from segregated to integrated states confers a computational alteration in the brain, which may be advantageous for certain cognitive tasks [18]. We thus reinterpret the gain-mediated transition in the functional configuration of the network in terms of the effective influence that neural regions can have over one another within a complex, adaptive, dynamical system [5]. Namely, subtle alterations in the neural gain control parameter lead to large transitions within the state space of functional topology, even within the constraints imposed by a hard-wired structural scaffold, with the resulting modulation of information processing capacity of the brain represented in different patterns of neural effective connectivity [1].

**Fig 5.**
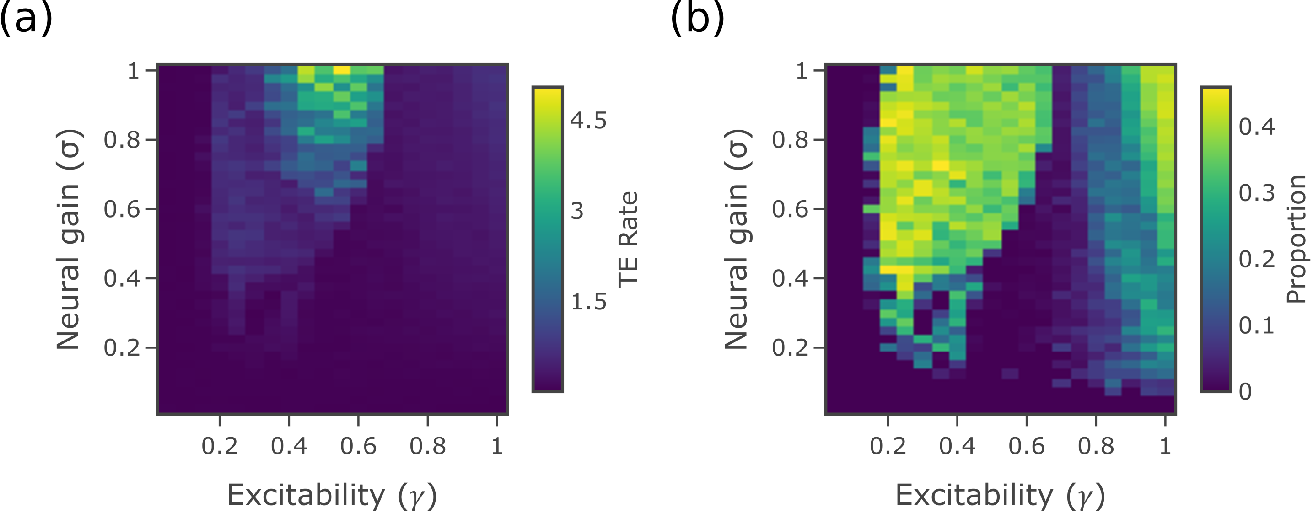
Information transfer between hemispheres. (a) TE rate over interhemisphere causal edges. (b) Proportion of significant TE rate occuring between interhemisphere source and target

In previous work [7], we identified a distinct boundary that was mediated by alterations in neural gain, which has long been linked to the functioning of the ascending (noradrenergic) arousal system [19]. Extending this previous work into the domain of information processing, we here observed a qualitative shift in regional computational capacity on either side of the gain-mediated phase transition. Namely, information storage (Fig. 2a) was maximal in the subcritical region (at intermediate γ), whereas information transfer (Fig. 3) peaked in the supercritical region. This result is strongly aligned with the observed transition in phase synchrony observed in previous related studies. In a system of Kuromoto oscillators, Ceguerra et al. [50] showed that the synchronization process can be modelled as a distributed computation, with larger and increasing transfer entropy associated with more strongly synchronized or integrated network states. In the case of our model, we expect that maintaining synchronisation in the face of noise requires strong ongoing transfer between the relevant regions.

Despite the strong qualitative effects observed at intermediate γ, the relationship between gain and information processing was distinctly non-monotonic. By tracking information-theoretic measures across the parameter space, we were able to distinguish six unique zones with qualitative differences in information processing dynamics (Fig. 6). For example, Zone 4 contained globally-synchronized oscillations which were relatively large, and also exhibited the strongest information transfer values. Note that this is not directly because the absolute range of the variables is larger (information measures on continuous-valued variables are scale independent [38]) but specifically due to variations in the relationships between dynamics of the regions. The differences between Zone 5 and Zone 6 – both of which occur at high γ but have distinct between-hemispheric TE (TE6 ≫ TE5, Fig. 5b) and AM rate (Fig. 2a, including different dependencies on local network structure and self-loops in Fig. 2e and f) – are also of interest, as they suggest that there may be distinct information processing signatures related to increasing multiplicative and response gain [52] to maximal levels, as in the case of epileptic seizures [53]. Future work should attempt to determine whether these categories are consistent across generative models, or perhaps relate to individual differences in topological recruitment across diverse cognitive tasks [17].

**Fig 6.**
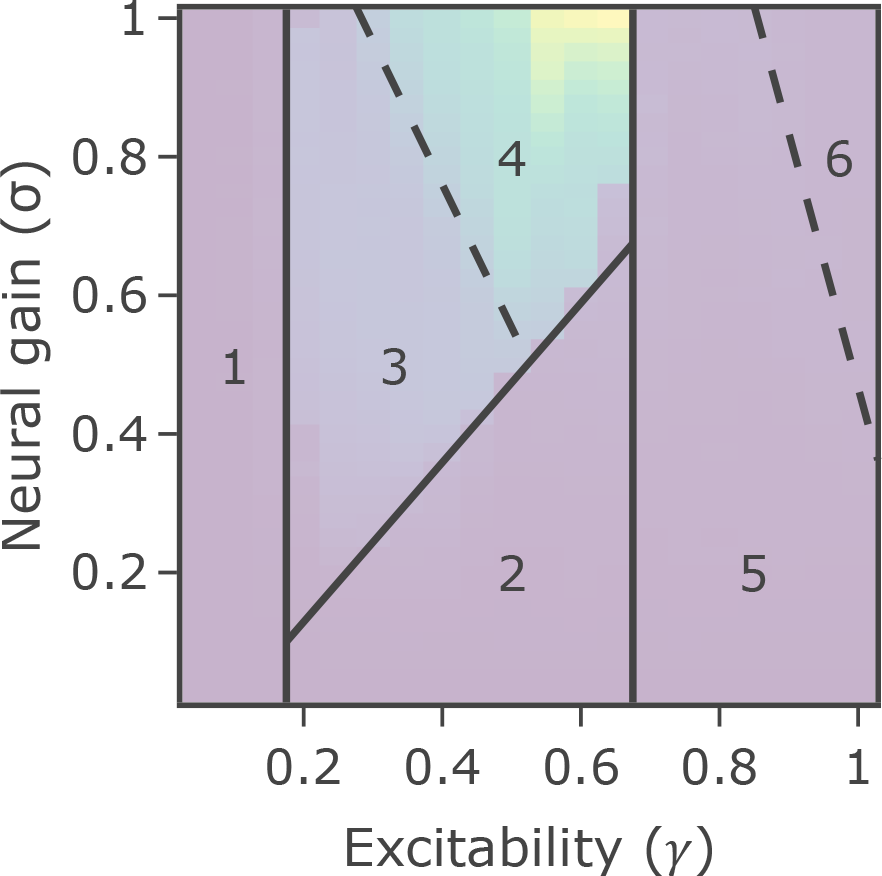
Phase portrait showing six identified regions. A transparent figure of the TE rate from Fig. 3a is shown behind for comparison. Dotted lines represent a looser boundary, which are not observed in all measures

As a general framework for understanding distributed computation within complex systems, the translation of the previous results into the language of information storage and information transfer allows their comparison to other systems whose information dynamics have been shown to undergo phase transitions, including artificial neural networks [31], random Boolean networks [35, 36] as models for gene regulatory networks, the Ising model [33] and indeed Kuramoto oscillators [50] as mentioned above. There appears to be substantial universality among the results from these systems, with similar patterns of information storage and transfer often observed around critical phase transitions – and crucially these patterns are echoed here in transitions driven by alterations to neural gain parameters in our neural mass model. Across all of these systems, we consistently observe that dynamics of subcritical states are dominated by information storage operations underpinning higher segregation, whilst information transfer amongst the components of the networks plays a much more significant role in the dynamics of supercritical states leading to higher integration. In contrast to both, the dynamics of the critical state exhibit a balance between these operations of information storage and transfer – a result which we emphasize was specifically observed again for the neural gain driven transitions examined here.

These insights allow us to address the question posed in our Introduction: are there computational advantages for the brain to operate in a near-critical state? In alignment with these results from other systems, and hypothesised as discussed earlier [13, 15, 16], the balance between these operations exhibited near the critical state could be expected to support a wide range of general purpose cognitive tasks (requiring both types of operations), as well as in allowing flexibility for rapid transitions to either sub- or supercritical behaviour in order to alter the computational structure and dynamics as required. Indeed it is straightforward to identify situations that would require rapid transitions away from criticality toward more segregated or integrated operation. A relatively segregated, modular architecture is comprised of regions with high information storage, suggesting that situations in which a more segregated architecture is beneficial to cognitive performance – such as a motor-learning task [54] or visual vigilance [55] – may retain their capacity for improved performance by promoting heightened information storage. In contrast, cognitive states associated with integration–such as working memory [17] or attention [56, 57] – may reflect heightened inter-regional influence, and hence, information transfer between the diverse specialist regions housed within distinct locations in the cortex and subcortex [18]. The flexibility inherent in operating near a critical state would be crucial in supporting rapid transitions to support either broad type of task.

The above interpretations align with a broader conjecture regarding utility of critical dynamics, such as in the “edge of chaos” hypothesis [28, 34] as well as more specific considerations regarding the utility of operating near criticality (but not directly at the “edge of chaos”) for the brain [13]. This convergence of results across the aforementioned systems suggests that the rules governing the organisation of whole brain dynamics may share crucial homology with other complex systems, in both biology and physics. However, inferring direct algorithmic correspondence will require more focused, direct comparison between the different systems. Furthermore, work remains to explain conditions leading to subtle differences in the patterns exhibited across the systems, for example the additional *maximization* of information storage and transfer capabilities near criticality in some transitions (e.g. Ising model [33]) but only a crossover without maximization in others (such as Kuramoto model [50] and the neural dynamics here).

The approach of information dynamics also provides a computational description of the dynamics of the system as they unfold at a local or point-wise level through time and across space [58]. Such descriptions provide quantitative insights regarding Marr’s “algorithmic” level [59] of how entities are represented within and operated on by a neural system [60]. In this study, we have not focused on the temporal dynamics of any particular task, but instead have examined the distribution of information processing signatures across the network. In particular, we have identified how the informational signature of brain dynamics relates to network structure as we transition across the neural gain parameter space. While the underlying network structure does not change, we seek to understand how its impact on the dynamics varies across the parameter space. Our approach allowed us to tease apart the relative importance of local network-supported versus internal mechanisms for information storage (Fig. 2), where the local network support explained much of the storage (as suggested for different dynamics [45]) except for within the strongly segregated regime. We also found that source regions with large in-degrees tended to be stronger information transfer sources, again for much of the parameter space except the strongly segregated regime. This aligned with our hypotheses and findings in other systems [50, 51], as well as related results such as that the degree of a node is correlated to the ratio of (average) outgoing to incoming information transfer from/to it in various dynamical models (including Ising network dynamics on the human connectome) [61, 62]. Finally, we compared the information transfer between hemispheres with the information transfer within hemisphere. Large proportions of information transfer could be apparently observed between non-directly linked regions across hemispheres in the supercritical regime, suggesting that the relatively large information transfer on the small number of inter-hemispheric causal edges supports significant global integration in this regime.

These local views of network structure were thus linked to whole-brain macroscopic topology in important ways. The extent to which the patterns triggered by changes in neural gain are targeted or global is a crucial question for future research, particularly given the recent appreciation of the heterogeneity of firing patterns within the locus coeruleus [63].

We note that the measures of information processing were estimated here using a Gaussian model, assuming linear interactions between the variables. This estimator was selected for efficient performance on the large data set. Although such estimators may not directly capture strongly non-linear components of the interactions, they nevertheless provide a useful descriptive statistic even when the linear-Gaussian model is violated. We note that the linear component often dominates (e.g. [64]), and indeed the larger embeddings such estimators support provide additional terms to indirectly model non-linear components in AIS and TE.

The motivation for the previous study [7] was an attempt to explain the mechanistic basis of fluctuations in functional network topology that were observed in empirical BOLD data [17], which were hypothesized to be functionally related to ongoing dynamics in the ascending arousal system [17, 65]. However, the sluggish temporal nature of the haemodynamic response typically clouds the interpretation of causal or indeed effective connectivity between brain signals [66, 67]. In particular, variable delays between neural activity and peak haemodynamic response around the brain means that temporal precedence in the BOLD response does not necessarily imply neuronal causality. While approaches have been suggested to address this issue [68, 69], we instead investigated information-theoretic signatures on simulated neural data, which has a much higher effective sampling frequency than BOLD, and is also relatively unaffected by the temporal convolution that masks neural activity in the BOLD response. In doing so, we highlight important multi-level organisation within the simulated neural time series, in which whole-brain topological signatures (measured using BOLD) overlap with specific signatures of regional (neural) effective connectivity. It remains an open question whether this relationship holds in empirical data; Increasing availability of intracranial human sEEG data will allow this to be tested more directly than with BOLD. In any case, our approach certainly holds promise for advancing our interpretation of fluctuations in global network topology across cognitive states [18, 70, 71].

In conclusion, we have shown that modulating neural gain in a biophysical model of brain dynamics leads to a shift in the computational signature of regional brain activity, in which the system shifts from a state dominated by self-referential information storage to one distinguished by significant inter-regional effective connectivity. These results provide a crucial algorithmic foundation for understanding the computational advantage of whole-brain network topological states, while simultaneously providing a plausible biological mechanism through which these changes could be instantiated in the brain – namely, alterations in the influence of the ascending arousal system over inter-regional connectivity.

## Materials and methods

### Simulation of Neural Activity

Neural activity was modelled (as per [7]) as a directed network of brain regions, with each region represented by an oscillating 2-dimensional neural mass model [41] derived by mode decomposition from the Fitzhugh-Nagumo single neuron model [72]. Directed coupling between 76 regions was derived from the CoCoMac connectome [73] with axonal time delays between regions computed from the length of fiber tracts estimated by diffusion spectrum imaging [43]. The model was simulated by stochastic Heun integration [74] using the open source framework The Virtual Brain [43].

The neural mass model at each region is given by the Langevin equations 1 and 2, which express the dynamics of local mean membrane potential (*V*) and the slow recovery variable (*W*) at each regional node *i*:

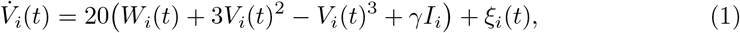

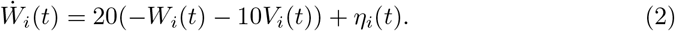

Here, all *ξ*_*i*_ and *η*_*i*_ are independent standard Wiener noises, and *I* is the synaptic current, given by

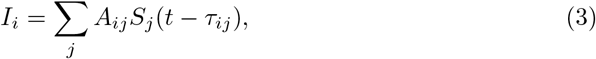

where *A*_*ij*_ is the connection weight from *j* to *i* in the directed connectivity matrix and *τ*_*ij*_ is the corresponding time delay from *j* to *i*. A sigmoid activation function was used to convert membrane potentials to normalized firing rates *S*_*i*_, where *m* = 1.5 was chosen to align the sigmoid with its typical input:

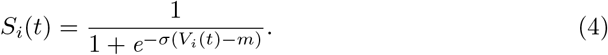

Using this model, we modulate the inter-regional coupling by varying the parameters for gain (*σ* in (4)) and excitability (γ in (1)) over a range of values between 0 and 1. At each parameter combination, membrane voltage (*V*_*i*_(*t*)) over time for each region was recorded as the time series input for the analysis of information dynamics. A sample length of 100,000 values per time series was used in the following analysis, corresponding to 50 seconds of one sample per 500 microseconds. The 2 kHz sampling rate was selected as described regarding the transfer entropy measure below. Each iteration was started from a different random initial condition.

Code implementing the model is freely available at https://github.com/macshine/gain_topology [75].

### Measures of Information Dynamics

The framework of information dynamics uses information theoretic measures built on Shannon entropy to model the storage, transfer and modification of information within complex systems. It considers how the information in a variable *X*_*n*+1_ at time *n* + 1 can be modelled as being computed from samples of this and other processes at previous times. Information modelled as being contributed from the past of process *X* is labelled as information storage, while information modelled as contributed from other source processes *Y* is interpreted as information transfer.

The Java Information Dynamics Toolkit (JIDT) [76] was used to calculate these measures empirically using the time series of neuronal membrane voltage from the 76 regions. For each combination of *σ* and γ parameter values, the active memory rate was calculated for each region, and the transfer entropy rate was calculated for each combination of two regions. Collective and conditional transfer entropy rates were also calculated. Each of these measures is explained in the following sections. The linear-Gaussian estimator in JIDT was utilized in these calculations (which models the underlying processes as multivariate Gaussians with linear coupling). As per our Discussion, this remains a useful descriptive statistic even when the assumed model is violated.

#### Active Information Storage

Active Information Storage (AIS) [39] models the contribution of information storage to the dynamic state updates of a process *X* by measuring how much information from the past of *X* is observed in its next observation *X*_*n*+1_. It is defined as the expected mutual information between realizations 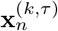 of the past state 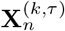 at time *n* and the corresponding realizations *x*_*n*+1_ of the next value *X*_*n*+1_ of process X [39]:

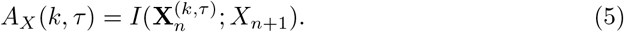

Formally, the states 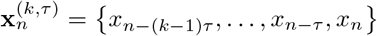 are Takens’ embedding vectors [77] with embedding dimension *k* and embedding delay *τ*, which capture the underlying state of the process *X* for Markov processes of order *k*. In general, an embedding delay of *τ* ≥ 1 can be used, which may help to better empirically capture the state from a finite sample size. (Note that non-uniform embeddings can be used [78]).

The determination of these embedding parameters followed the method of Garland et al. [79] finding the values which maximize the AIS, with the important additional inclusion of bias correction (because increasing *k* generally serves to increase bias of the estimate) [80]. For several sample *σ*, γ pairs in both the sub- and supercritical regimes we examined these parameter choices across all regions (up to *k*, *τ* ≤ 30), and found the optimal choices to be consistently close to *k* = 25 and *τ* = 12 for all variables (for the sampling interval Δ*t* = 0.5 ms). As such, *k* = 25 and *τ* = 12 were used for all investigations.

Note that while a larger AIS is likely to give rise to a larger auto-correlation time, there are significant differences between the two which make AIS a much more powerful measure, and directly relevant for modelling the utilisation of information storage (unlike autocorrelation). Primarily these differences stem from AIS examining the relationship between multiple past values (as the embedded past) to the current value of the time series, taking into consideration whether those past values are providing the same information redundantly or unique information, or indeed are synergistically providing more when they are considered together. Auto-correlation values in contrast only ever examine relationships from one past value to the current, and are unable to resolve such complexities in the process. (As an information-theoretic measure, AIS can also capture non-linear interactions, although in this study we only use a linear estimator). This leads to the AIS providing very different values to auto-correlation, and indeed much richer insights. For example, significant reductions were observed in AIS in multiple regions of ASD subjects versus controls [81], indicating significantly reduced use or precision of priors in dynamic state updates of ASD subjects. In contrast, no such differences were observable using auto-correlation times or signal power.

Because of the fast sample rate (Δ*t* = 0.5 ms) of the neuronal time series, proper analysis requires using a formulation of the information theoretic measures suitable for continuous time processes. In general, this means that information storage and information transfer are conceptualized as measures that *accumulate* over some finite time interval at an associated *rate* [44]. Both the accumulated and rate measures, however, diverge in the limit as the time step approaches zero. Intuitively, for continuous processes such as those here, this is because all information about the next time step can be captured by the previous time step in the limit as the two samples become essentially identical. These divergent properties can be circumvented by decomposing active information storage into components, 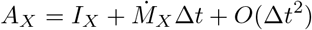, comprising [44]:

- the instantaneous predictive capacity which measures the information storage from the immediately previous time step, *I*_*X*_ = *I*(*X*_*n*_; *X*_*n*+1_), and
- the **active memory utilisation rate**(AM rate) which measures the additional accumulation rate of information storage from time steps before that, 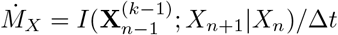.

The instantaneous predictive capacity inherits the divergent nature of the active information storage, while the active memory utilization rate takes on the intuitive representation of memory as a rate [44]. Crucially, 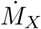 converges to a limiting value as Δ*t* → 0 for well-behaved continuous processes such as those considered here unlike *A*_*X*_ and *I*_*X*_ (see full details in [44]), and thus only 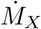 is used in our investigations here. As a rate, the units of measurement of 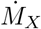 are in bits per second. As recommended by Spinney et al. [44, 46], Δ*t* = 0.5 ms was selected on confirming that the transfer entropy rate (see next subsection) and 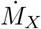 are stable to Δ*t* in this regime and appear to have converged to a limiting value as Δ*t* → 0.

#### Transfer Entropy

Transfer entropy [40, 82] models the contribution of information transfer from a source process *Y* to the dynamic state updates of a destination (or target) process *X* by measuring the amount of information that *Y* provides about the next state of process *X* in the context of the destination’s past. This perspective of modelling of information transfer after first considering storage from the past contrasts the two operations, and ensures that no information storage is attributed as having been transferred [82].

Quantitatively, this is the expected mutual information from realizations 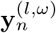 of the state 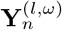 of a source process *Y* to the corresponding realizations *x*_*n*+1_ of the next value *X*_*n*+1_ of the destination process *X*, conditioned on realizations 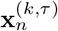 of its previous state 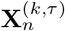:

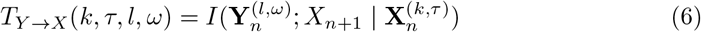

In general, an embedding of the source state 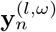 with *l* > 1 could be considered as this would allow *Y* to be a Markovian process where multiple past values of *Y* in addition to *y*_*n*_ are information sources to *x*_*n*+1_. However for this analysis only *l* = 1 previous time step of the source process is used (denoted *T*_*Y* →*X*_ (*k*, *τ*)), in line with the known function of the neural model in (1)-(4).

In order to best model information transfer, the source processes for TE measurements are constrained to the known causal information contributors [47]. In this case these are the upstream parents of the target in the structural connectivity matrix.

As mentioned in the previous subsection, the small time steps of the neuronal time series requires us to consider continuous-time formulations, meaning we compute a **transfer entropy rate**, 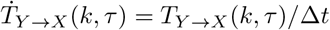 [44, 46].

The use of the linear-Gaussian estimator in JIDT for TE estimation makes the calculated transfer entropy (rate) equivalent, up to a constant, to Granger causality (rate) [83, 84].

Finally, note that transfer entropy estimations can be non-zero even where the source and destination processes have no directional relationship, due to estimator variance and bias (see summary in [82, Sec. 4.5.1]). As such, one can make a statistical test of whether a transfer entropy estimate is statistically different from the null distribution of values that would be observed for source and destination processes with similar properties but no directed relationship. We perform a test of statistical significance for each directed pair of processes in producing Fig. 5b, retaining there only the transfer entropies for pairs that were determined to be statistically significant against the null distribution against a p-value threshold of *α* = 0.05. This test is carried out analytically for the Gaussian estimator, as described in [76, App. A.5], with a Bonferroni correction for all directed pairs that are tested.

#### Conditional and Collective Transfer Entropy

Higher order terms of information transfer can capture the multivariate effects from multiple sources to a single target. Two higher order terms which were calculated are the conditional and collective transfer entropies.

Conditional transfer entropy [48, 49, 85] extends the basic form of transfer entropy by conditioning on the history of another source process, *Z*. This captures the mutual information between the past of source *Y* and the next value of target *X*, conditioned on both the history 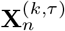 of *X* and the history of conditional source *Z*:

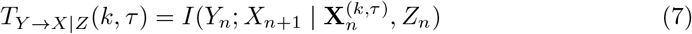

Of course, the above may in general incorporate embeddings for both *Y* and *Z*, and can be extended to condition on several other sources **Z** (excluding *Y*) at once. It should be noted that a conditioned transfer entropy can be either larger or smaller than the unconditioned measure, in the same way that a conditional mutual information can both increase due to the addition of synergistic information that can only be decoded with knowledge of both the source and conditional, as well as decrease due to a removal of redundant information provided by both the source and conditional. The conditional transfer entropy thus includes unique information from the source but not the conditional, and synergistic information provided by the source and conditional together, in the context of the past of the target [48, 86]. These components cannot be pulled apart using the tools of traditional information theory, but efforts are being made by approaches of Partial Information Decomposition (PID) [87–89].

At the same time, collective transfer entropy [49, 90] models the total information transfer from a set of sources to a target, capturing unique information from each source, avoiding double-counting redundant information across the sources, and capturing multivariate synergistic effects. Given a multivariate set of sources **Y**, this refers to the measure *T*_**Y**→*X*_ (*k*, *τ*) (again ignoring possible embeddings on the **Y** processes).

For these experiments, we compute conditional transfer entropy rate and collective transfer entropy rate, similar to the pairwise transfer entropy rate. Only the highest order conditional transfer entropy is calculated (known specifically as complete transfer entropy [48, 49]). This means that for each causal source, the conditional transfer entropy to a particular target involved conditioning on *all* the other causal sources to that target, as identified from the directed connectome. Also, we calculated the collective transfer entropy to a given target from all causal sources to that target region, as identified from the directed connectome.

### Network Motifs

The 76 regions of this model are connected by a directed, weighted network *A*_*ij*_ (see (3)) derived from the CoCoMac connectome [73]. The information storage of each region is expected to be related to the number of certain network motifs which provide feedforward and feedback loops involving that region. For linearly-coupled Gaussian processes the active information storage can be calculated as a function of weighted counts of these motifs [45]. However, for the more complex dynamics of this system, we cannot derive an exact relationship. Instead, we approximate how well we expect the network structure to support storage at a particular node or region *a* with a weighted linear combination of these motif counts. This **local network support** for memory is then correlated with the observed active memory rate (across all nodes for a given *σ*, γ) to see where in the parameter space this expectation held true.

The motifs which were considered (based on those identified in [45]) incorporated feedback loops including the target node *a* and feedforward loops terminating at node *a* (see Fig. 2d). The weighting given to each motif depends on the number of incoming links to *a* and their edge weights (which are derived from the coupling strengths *A*_*ij*_). First the edge weight of each link is normalized (taking inspiration from generation of normalized Laplacians [91]) by the total incoming edge weight for each target (excluding self loops) to generate *C* = *D*^−1^(*A* − diag(*A*_1,1_, *A*_2,2_,…, *A*_76,76_)), where *D* = diag(*d*_1_, *d*_2_…, *d*_76_) with *d*_*i*_ = ∑_*j*≠*i*_ *A*_*ij*_. The local network support Ψ_*a*_ at node *a* is then computed as a linear sum of the relevant motifs at node *a* using the normalized weights *C*:

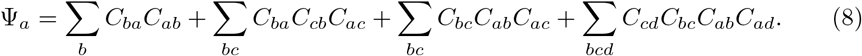

Note that the four weighted motif counts in the equation for Ψ_*a*_ correspond respectively to the four motifs shown in Fig. 2d to contribute to storage in the dynamics of node *a*. The contribution from longer motifs diminishes with length due to the normalisation, and so we limit (8) to the shortest two contributing feedback and feedfoward motifs (except for any self-loop at *a*).

Note that the self-loops were ignored in the local network support measure in order to consider relative support for memory from distributed network effects only in Ψ_*a*_. The relative contribution of self-loops *A*_*ii*_ (i.e. synaptic connections between neurons within the same brain region) to memory was instead analysed separately. A similar weighting was applied to self loops in order to normalize their contribution with respect to total incoming edge weights (this time including the self loop). Here, we first computed *F* = *G*^−1^*A*, where *G* = diag(*g*_1_, *g*_2_…, *g*_76_), *g*_*i*_ =∑_*j*_ *A*_*ij*_. Then, in order to evaluate the relative strength of contribution of self-loops to memory across the parameter space, we correlated the *F*_*ii*_ with the observed active memory rate across all nodes for each given *σ*, γ pair. Note that we focus here on the synaptic connections between neurons within the same brain region modelled by *A*_*ii*_, which is mediated by the neural gain parameters. This analysis does not include the feedback terms in *V* and *W* in (1) and (2) which correspond to self-coupling in the oscillatory dynamics of individual neurons that make up the population. We do not include those terms because they are constant across regions and are not moderated by the neural gain parameters.

## Acknowledgments

MJA was supported through a Queensland Government Advance Queensland Innovation Partnership grant AQIP12316-17RD2. JL was supported through the Australian Research Council DECRA grant DE160100630. JMS was supported through a University of Sydney Robinson Fellowship and NHMRC Project Grant 1156536. JMS and JL were supported through The University of Sydney Research Accelerator (SOAR) Fellowship program. High performance computing facilities provided by QIMR Berghofer Medical Research Institute and The University of Sydney (artemis) have contributed to the research results reported within this paper. The funders had no role in study design, data collection and analysis, decision to publish, or preparation of the manuscript.

